# Notes on the diet composition of Anoles lizards, *Anolis* (Dactyloidae), in the Yasuni National Park (Ecuador)

**DOI:** 10.1101/2023.06.15.545064

**Authors:** Javier Pinto

## Abstract

In this study, we observed and briefly described the diet of six species of lizards of the genus Anolis in the Yasuni National Park, located in the western part of the Amazon Rainforest. A total of 241 items found in the stomachs of the lizards were classified. We noted that Aranea and Hymenoptera were the most frequent diet categories used by the lizard community. In terms of volume, Hemiptera and insect larvae were the most representative prey. We aim that the description of the diet of *Anolis* provided by this study can be further combined with information pertaining to their natural history, thus shedding light on ecological mechanisms that influence adaptation.

## Introduction

The genus *Anolis* (Iguanidae: Dactyloinae) is a widely distributed group in the Caribbean, Central America, and northern South America (Losos, 1992; Losos & Ricklefs, 2009) and represents the most diverse genus of lizards on the planet (Uetz & Hošek, 2022). Despite being extensively studied in the Caribbean, information on Anoles on the continent is scarce. On the mainland, *Anolis* lizard communities are usually part of very large communities of insectivorous lizards. Diet constitutes a pivotal part of reptile’s natural history, due it provides energy to perform biological processes such as growth and reproduction (Dunham *et al*., 1989). Previous studies had described *Anolis* diet in several locations within the Amazon Rain Forest (Vitt et al., 2002, 2003a, 2003b). This set of data describes the diet of a community of *Anolis* in one of the most biodiverse places on the planet located in the Western Amazon Forest, the Yasuní National Park (Bass et al., 2010), consisting of six species: *A. fuscoauratus, A. ortonii, A. scypheus, A. transversalis, A. trachyderma*, and *A. punctatus*.

*A. fuscoauratus* forages in the lower strata of the forest, it is a generalist although some populations show a preference for the Orthoptera (Vitt et al., 2003a; Vitt & De la Torre, 1996). *A. ortonii* and *A. scypheus* are passive foragers, seeking for food in leaf litter and low vegetation (Vitt & De la Torre, 1996; Duellman, 1978). *A. trachyderma* is also passive, semi-arboreal forager that also feeds on invertebrates in leaf litter. *A. transversalis* and *A. punctatus* are species that mostly inhabit the canopy (Vitt et al., 2003b). The description of diet of *Anolis* provided by this study can be further combined with information pertaining to their natural history, thus shedding light on ecological mechanisms that influence adaption.

## Material and Methods

### Lizard sampling

Sampling was carried out for 70 days during June, July, August, and September 2012 in Yasuní National Park, Orellana Province, Ecuador, on the southern bank of the Tiputini River (76° 24’
s 1.8 ′′ W; 0′ 40′ 16.7′′ S).

### Specimen capture and stomach content obtention

*Anolis* Individuals were searched in different forest strata using binoculars. During the sampling months, 112 individuals were observed, 91 were captured and the stomach content obtained. Once the lizard was captured, stomach content was extracted in the laboratory. A ball-tipped dosing needle was introduced through the mouth to the level of the stomach. Using a syringe attached to the needle, water was injected, and the needle was moved back and forth while the stomach was massaged until the lizard regurgitated. The obtained items were preserved in 95% alcohol for subsequent analysis. Once all stomach contents were obtained, each prey was identified with the help of a stereoscope using taxonomic keys for insects and arachnids (Tripplehorn & Johnson, 2005). Identification was performed at the taxonomic level of order for most of the preys and some to level of family. After the stomach content was obtained, the lizard was fed with small crickets and released back into the place of capture.

After identification, the stomach contents were storage in Eppendorf tubes containing 95% ethanol for preservation. The tubes are storage in the Zoology Museum of Catholic University in Quito, Ecuador.

### Data analyses

Once the prey consumed was identified, the amount of prey ingested by each lizard was determined, and thus, the diets of each species could be categorized. The volume of each individual prey was calculated using the spheroid formula: V = 4/3Π (length/2) (width/2)2. In one category (no identifiable), we classified prey that could not be identified due to various factors, for example, the state of digestion of the prey. Plant tissue was found in three lizards, whereas lizard scales were found in four individuals, and empty stomachs were found in eight individuals.

## Results and Discussion

Among identified prey, Aranea was the most abundant category (n= 37), followed by Hymenoptera (n= 33), Hemiptera (n= 32), Orthoptera (n= 28), and insect larvae (n= 24). The least frequent consumed categories were: Collembola (n= 2), insect exuviae (n= 2), Isoptera (n= 2), Blattodea (n= 1) and Phasmidae (n= 1). Considering the volume of consumed prey, the most used category was Hemiptera (28.41%), followed by insect larvae (22.86%). Other categories consumed considerably were Orthoptera (12.02%), Aranea (11.76%), Coleoptera (10.53%) and Hymenoptera (7.05%), while the least consumed categories in term of volume were Blattodea, Collembola, Diplopoda, Isopoda, Isoptera, Phasmidae and exuviae. Eight lizard had empty stomachs and 16 out of 241 items were not identified.

At lizard species level, a total of 115 items were found in the stomachs of *A. fuscoauratus*, mostly represented by Aranea (n= 23 and 29.69% of the total volume consumed), Hemiptera (n= 12 and 22.22% of the volume) and insect larvae (n= 11 and 11.63% of the volume). Several small Orthoptera were found (n= 14) but their volume represented only 6.05%. A small amount of plant tissue was found in one individual.

Thirty prey items were found in the stomachs of *A. ortonii*, Hymenoptera was the most consumed category (n= 8), represented 6.63% of the total volume consumed. Hemiptera (n= 6) represents 35.99% of the volume. A total of 26 prey items were found in the stomach contents of *A. scypheus*, insect larvae being the most representative category (n= 7 and 61.56% of the total volume). In one individual of *A. scypheus*, 19 Arthropoda eggs were found and their volume only represents 1.52% of the total volume consumed (not reported in Table 1). In *A. transversalis*, 43 prey were found, Hymenoptera was the most representative category (n= 11), although it only represents 7.35% of the total volume, followed by Hemiptera (n= 10 and 42.13% of the volume). A considerable amount of Coleoptera (n= 6 and 20.19%) and Aranea (n= 6 and 8.66%) were found. Although few larvae were found (n= 3), their volume represented 11.4% of their diet. In the case of *A. punctatus*, only seven prey items were obtained, mostly Orthoptera (n= 3 and 52.29% of the total volume) and Hemiptera (n= 2 and 46.96% of the total volume consumed). Similarly, due to the small number of individuals of *A. trachyderma* captured, only four preys were obtained, two Orthoptera (56% of the volume), and one insect larva (41.45% of the volume). A summary of the diet composition of the lizards captured is shown in Figure 1.

**Table 1.** Number and volume (mm^3^) of type preys consumed by *Anolis* lizards.

|  |  | *Anolis species* |  |  |  |  |  |  |  |
| --- | --- | --- | --- | --- | --- | --- | --- | --- | --- |
| No. | Item |  | *fuscoauratus* | *ortonii* | *transversalis* | *scypheus* | *trachyderma* | *punctatus* | Total |
| 1 | Araneae | Count | 23 | 5 | 6 | 2 | 1 | 0 | 37 |
|  |  | Volume | 774.79 | 33.82 | 272.15 | 12.85 | 2.69 | 0 | 1096.3 |
| 2 | Blattodea | Count | 0 | 0 | 0 | 1 | 0 | 0 | 1 |
|  |  | Volume | 0 | 0 | 0 | 33.79 | 0 | 0 | 33.79 |
| 3 | Coleoptera | Count | 17 | 4 | 6 | 2 | 0 | 1 | 30 |
|  |  | Volume | 251.12 | 17.91 | 634.53 | 76.14 | 0 | 2.08 | 981.78 |
| 5 | Collembola | Count | 1 | 0 | 1 | 0 | 0 | 0 | 2 |
|  |  | Volume | 6.65 | 0 | 10.98 | 0 | 0 | 0 | 17.63 |
| 6 | Diplopoda | Count | 2 | 0 | 0 | 1 | 0 | 0 | 3 |
|  |  | Volume | 13.08 | 0 | 0 | 19.31 | 0 | 0 | 32.39 |
| 7 | Diptera | Count | 3 | 4 | 1 | 0 | 0 | 0 | 8 |
|  |  | Volume | 72.87 | 61.11 | 24.08 | 0 | 0 | 0 | 158.06 |
| 8 | Exuviae | Count | 2 | 0 | 0 | 0 | 0 | 0 | 2 |
|  |  | Volume | 246.62 | 0 | 0 | 0 | 0 | 0 | 246.62 |
| 9 | Hemiptera | Count | 12 | 6 | 10 | 2 | 0 | 2 | 32 |
|  |  | Volume | 579.9 | 151.46 | 1324.21 | 228.57 | 0 | 365 | 2649.14 |
| 10 | Hymenoptera | Count | 9 | 8 | 11 | 4 | 0 | 1 | 33 |
|  |  | Volume | 142.62 | 27.9 | 231.04 | 252.47 | 0 | 3.74 | 657.77 |
| 11 | Insect eggs | Count | 1 | 0 | 0 | 1 | 0 | 0 | 2 |
|  |  | Volume | 0.28 | 0 | 0 | 33.58 | 0 | 0 | 33.86 |
| 12 | Isopoda | Count | 4 | 0 | 0 | 1 | 0 | 0 | 5 |
|  |  | Volume | 32.28 | 0 | 0 | 101.27 | 0 | 0 | 133.55 |
| 13 | Isoptera | Count | 2 | 0 | 0 | 0 | 0 | 0 | 2 |
|  |  | Volume | 6.42 | 0 | 0 | 0 | 0 | 0 | 6.42 |
| 14 | Larvae | Count | 11 | 2 | 3 | 7 | 1 | 0 | 24 |
|  |  | Volume | 303.55 | 41.09 | 358.34 | 1364.63 | 64.13 | 0 | 2131.74 |
| 15 | Lizard scales | Count | 3 | 0 | 0 | 1 | 0 | 0 | 4 |
|  |  | Volume | 1.24 | 0 | 0 | 0.41 | 0 | 0 | 1.65 |
| 17 | Orthoptera | Count | 14 | 1 | 5 | 3 | 2 | 3 | 28 |
|  |  | Volume | 157.93 | 87.57 | 287.82 | 93.22 | 87.91 | 406.38 | 1120.83 |
| 18 | Phasmatodea | Count | 1 | 0 | 0 | 0 | 0 | 0 | 1 |
|  |  | Volume | 19.23 | 0 | 0 | 0 | 0 | 0 | 19.23 |
| 19 | Vegetal tissue | Count | 2 | 0 | 0 | 1 | 0 | 0 | 3 |
|  |  | Volume | 0.38 | 0 | 0 | 0.41 | 0 | 0 | 0.79 |
|  | Total | Count | 115 | 30 | 43 | 26 | 4 | 7 | 241 |
|  |  | Volume | 2608.96 | 420.86 | 3143.15 | 2216.65 | 154.73 | 777.2 | 9321.55 |

**Figure 1.**
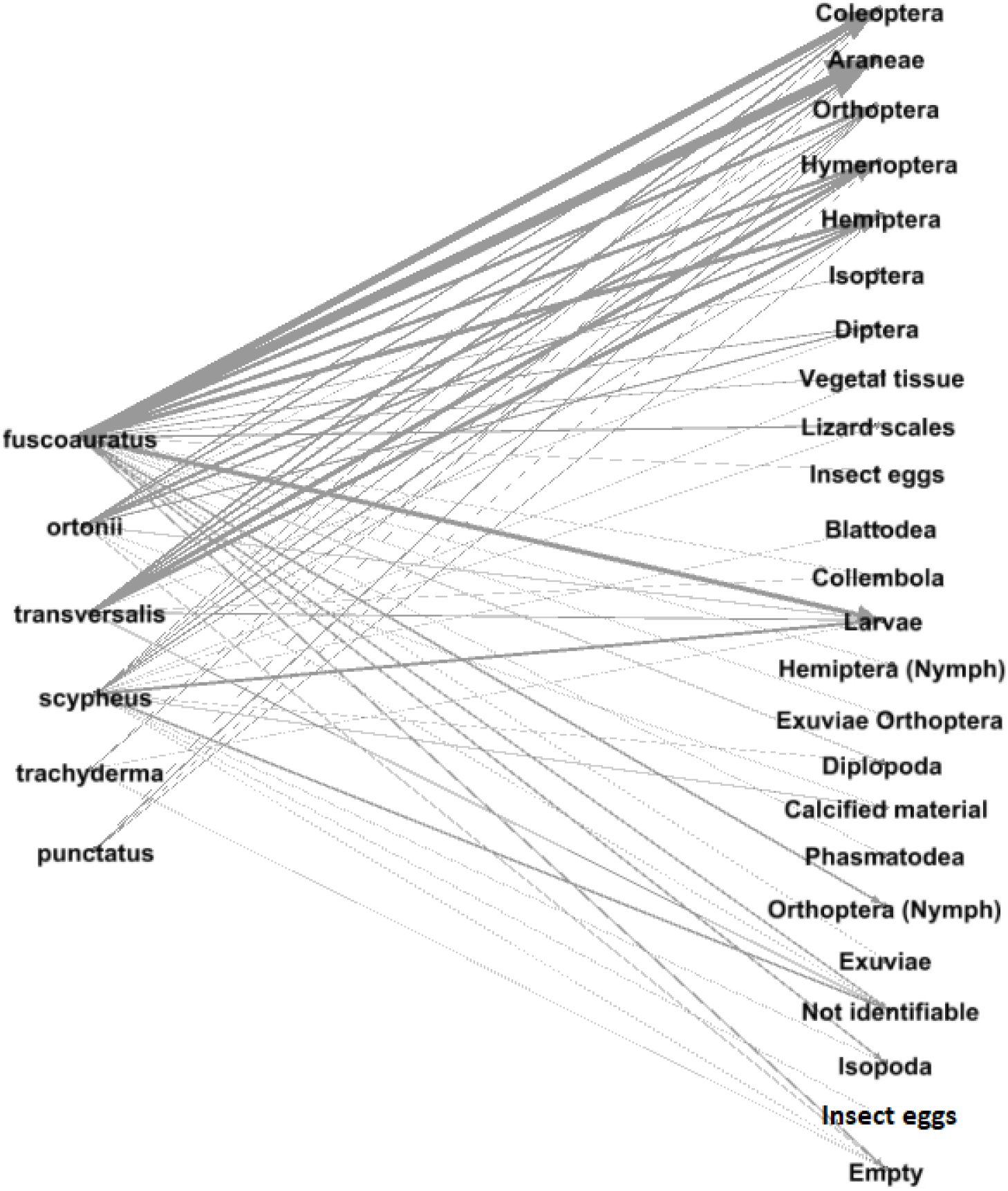
Bipartite network *Anolis* lizard-prey interaction from Yasuní National Park. The numbers of prey items found in stomach content are shown as links (grey arrows) connecting *Anolis* species and prey items. The left column contains the six species of *Anolis* lizards comprehended in this study, the right column includes all categories of lizards’ diet.

The species with the highest number of prey items observed per individual was *A. transversalis*, with an average of 3.83 prey items per lizard, followed by *A. punctatus* (2.33), *A. fuscoauratus* (1.93), *A. ortonii* (1.82), *A. scypheus* (1.73) and *A. trachyderma* (1.33). The difference between *Anolis* species in quantity and volume of prey consumed per individual was not significant (p= 0.565 and 0.769, respectively) Table 2. Detailed database is showed in Supplementary table 1.

**Table 2.** Prey number and dimensions consumed per lizard species. SD= Standard deviation.

| species | n | # Item per lizard | Media (item length) | SD (item length) | Media (item width) | SD (item width) |
| --- | --- | --- | --- | --- | --- | --- |
| *A. fuscoauratus* | 111 | 1.93 | 6.46 | 3.39 | 2.28 | 1.29 |
| *A. ortonii* | 32 | 1.82 | 5.73 | 3.58 | 1.75 | 0.83 |
| *A. transversalis* | 46 | 3.83 | 10.33 | 4.46 | 3.16 | 1.46 |
| *A. scypheus* | 33 | 1.72 | 9.6 | 5.75 | 3.19 | 2.22 |
| *A. trachyderma* | 4 | 1.33 | 9.19 | 4.42 | 2.55 | 0.93 |
| *A. punctatus* | 7 | 2.33 | 13.33 | 7.61 | 3.14 | 1.62 |

## Acknowledgements

Dr. Omar Torres-Carvajal, Dr. Andrés Mármol, the Staff of Yasuní Research Station of The Universidad Católica del Ecuador. This project was financed by Universidad Católica del Ecuador.

